# Structure-driven effects on genomic DNA damage propensity at G-quadruplex sites

**DOI:** 10.1101/2021.12.02.471014

**Authors:** Adib A. Abdullah, Claudia Feng, Patrick Pflughaupt, Aleksandr B. Sahakyan

## Abstract

Our genome contains about half a million sites capable of forming G-quadruplex (G4) structures. Such structural formations, often localised at important regulatory loci, have high capability of altering the predisposition of corresponding genomic spans to endogenous and exogenous DNA damage. In this work, we devised an approach to systematically enrich and zoom onto structure-driven effects on the propensity to undergo 9 types of DNA damage: ultraviolet radiation-induced pyrimidine-pyrimidone (6-4) photoproduct PP and cyclobutane pyrimidine dimer CPD couplings (two dyad-based subtypes in each), cisplatin-mediated G-G crosslinks, reactive oxygen species induced 8-oxoguanine damage, DNA fragmentation upon natural decay and fossilisation, breakages from artificial enzymatic cleavage and ultrasound sonication. Our results indicate that the structural effects on DNA damageability at G4 sites are not a simple combination of shielding (G4 strand) and de-shielding (opposite strand) against damaging factors, and the outcomes have different patterns and variation from one damage type to another, highly dependent on the G4 strength and relative strand localisation. The results are accompanied by electronic structure calculations, detailed structural parallels and considerations.

**Graphical Abstract**. An illustration of the considered damage factors acting on G-quadruplex sites, through strand-specific (two pointing thunder signs) and strand-invariant (a single pointing thunder sign) manner.

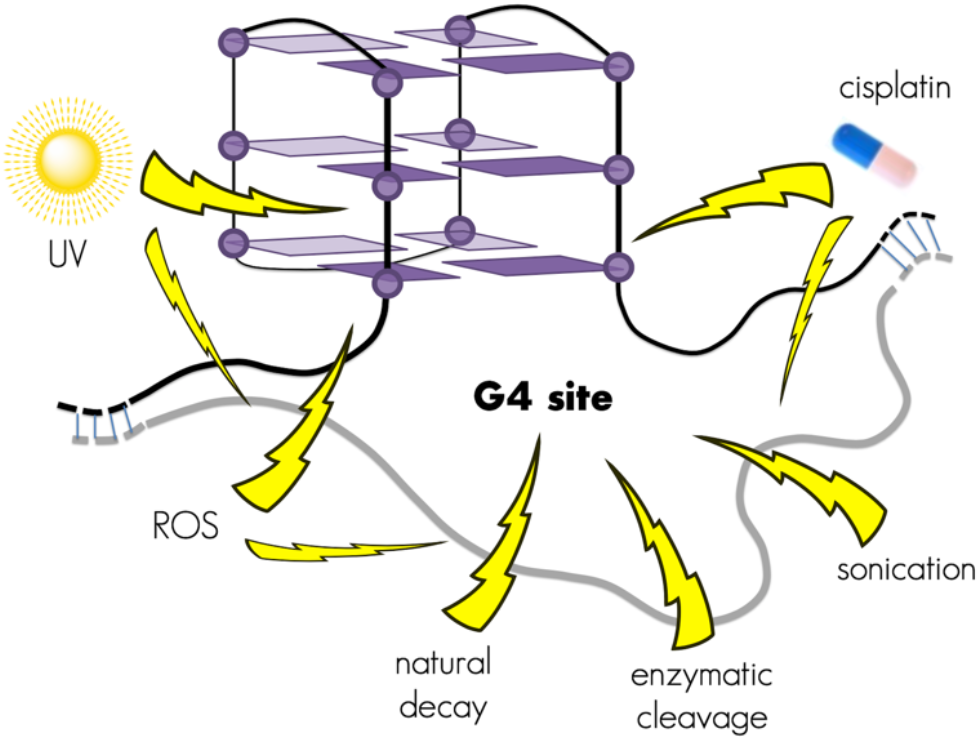

## INTRODUCTION

G-quadruplexes (G4s) are four-stranded DNA formations comprised of G-tetrads with the co-planar arrangement of guanine bases through Hoogsteen base pairing (**Figure 1A**), which are then stacked together to form G4s (**Figure 1B**) (1-3). Intramolecular (intra-strand) G4s, forming on a single strand of a DNA, are possible owing to the presence of four continuous G-tracts of guanine bases, interspersed with three loops that can each contain any base (**Figure 1B**). Such formations, about half a million in quantity within the human genome (4), leave the opposite strand into a more exposed single-stranded ssDNA state (**Figure 1C**), often open for other molecular interactions (5, 6) and, at times, capable of forming cytosine-driven i-motifs (7-9). In G4 sites, both DNA strands thus undergo influential re-organisations with much capability to modulate the exposure of those strands to various endogenous and exogenous damaging agents (10).

**Figure 1.**
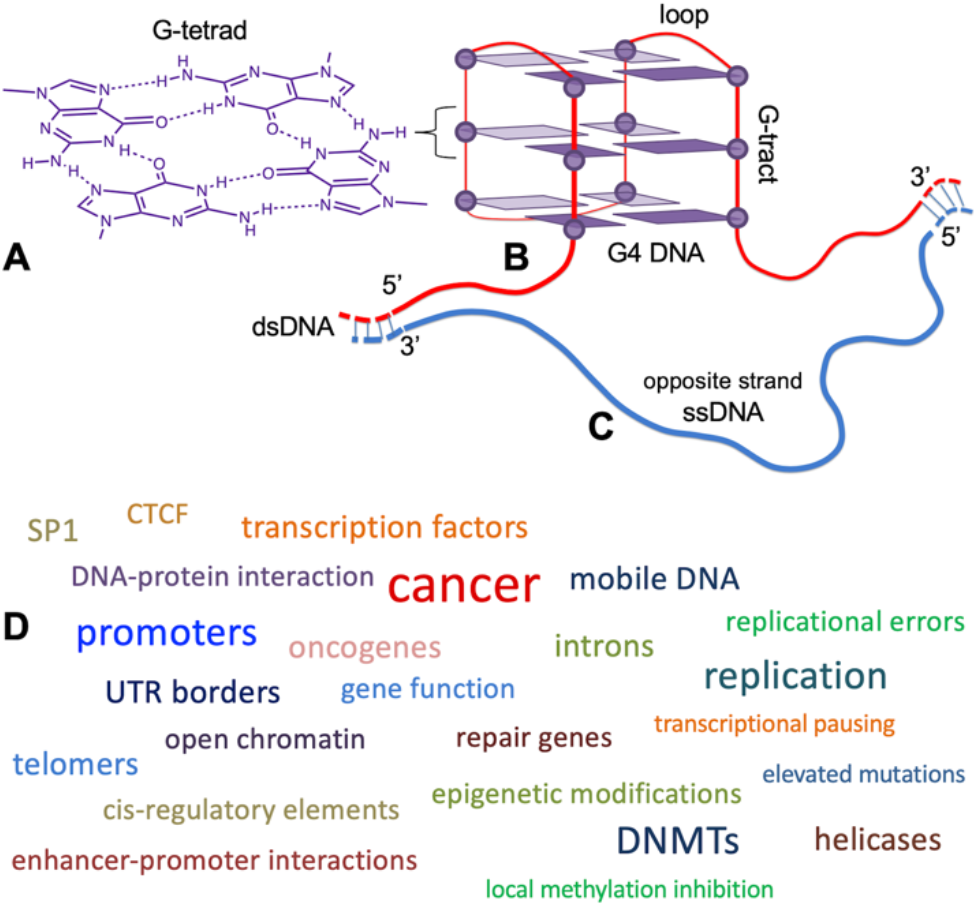
Structural characteristics of a G-quadruplex (G4) site in a genome. Schematic representation of an intra-strand G4 structure (**A**-**C**) is shown, along with the chemical structure of its constituent G-tetrad planes (**A**). Structural features of the G4 containing strand (**B**) and the opposite exposed strand (**C**) are shown. The major reported functional and genomic associations of G4 structures are highlighted in (**D**), with the brought terms hierarchically and non-linearly sized according to the number of Google search hits (November 2021) with the corresponding term and G-quadruplex.

In past, multiple studies focused on the influence of different epigenetic modifications (11, 12) and DNA lesions (13) on the formation and stability of G-quadruplexes, heralding the possibility of fine interplay between DNA fold and such modifications (10). Burrows and co-workers elucidated the role of the oxidative damage of guanine on non-B DNA structure formation, showing that those can have an impact switching on and off the genes through the stabilisation and destabilisation of co-localised structures (13, 14).

The aim of this study is to systematically investigate the influence of the structural factors at G4 sites on the predisposition of DNA to various types of damage. Our study was designed in a specific manner to zoom onto only the structure-driven effects, enriching it from the possible sequence effects and nucleobase frequency biases. This was done by grouping human genomic G4s in accordance with their stability categories, with further analyses of the damage propensity at G4 sites when G4 stability gradually increases. Any coupled gradual change in our damageability data would thus represent the effects of the increasingly structured G4 and unstructured opposite strands.

G4 structures have been found in abundance within our genome *in silico* (15, 16), *in vitro* (4) and in living cells (17, 18), and were shown to play variety of roles in gene regulation and genome organisation (2, 3, 19, 20). G4s were found to be particularly influential (**Figure 1D**) at around the important regulatory loci in our genome, such as at promoter sites (21-24) of genes, marking the borders of UTRs (25) and introns (26), serving as important transcription factor binding hubs (27-30), important in human telomers (31-34), present at mobile DNA elements (35) with evolutionarily modulated role in LINE 1 mobility (36), acting as DNA methyltransferase binding partners (37-39) with important mechanistic implication in DNA methylation patterns (40), present at important DNA repair genes (41) and oncogenes (42, 43), acting as *cis*-regulatory elements (44), influencing mutations (45, 46) and being an active target for many proteins (47) and helicases in particular (48). It is thus of paramount importance to account for G-quadruplex formation as an important factor while studying the influence of various agents on important regulatory spans of our genomic DNA.

This work reveals a well-defined and, where applicable, strand-specific dependence of susceptibilities to 9 types of DNA damage and fragmentation. Accompanied by a wealth of structural parallels and electronic calculations, we highlight the structural effects on DNA damageability at G4 sites going beyond the expected combination of shielding (G4 strand) and de-shielding (opposite strand) against damaging factors. Our outcomes have different patterns and variation from one damage type to another, highly dependent on the G4 strength and relative strand localisation.

## MATERIALS AND METHODS

### General notes on the performed calculations

The developed workflows and analyses in this study employed the R programming language (49). The resource demanding computations were performed on a local Linux-based computing cluster at MRC WIMM, University of Oxford, by using nodes with 3 × 2.7 GHz 8-core E5-2680 Intel Xeon processors and 256 GB random access memory. Analyses involving the human genome were performed using the unmasked version of the human reference sequence hg19/GRCh37, as accessed from Ensembl (50) genome database.

### Processing of genomic G-quadruplex data

Data from the G4-seq study were taken from (4), as deposited on GEO (51) under the accession identifier GSE63874. We used the dataset corresponding to the potassium ion treatment in the sequencing buffer, as opposed to the one utilising G-quadruplex-stabilising non-physiological ligand. G4-seq data presents a genome-wide profile of base mismatch levels (mm%) caused by polymerase stalling upon G-quadruplex (G4) encounter in sequencing. The experimental mm% values are descriptive of the stability of a G4 residing near the mm% measurement site, with high mm% reflecting higher stability of the formed G4 structure (52). To assign unique mm% values per putative quadruplex sequence (PQS), the procedure was mirrored from (52). In particular, the human genome was scanned through the extended PQS motif with [G_3+_N_1−12_]_3+_G_3+_ regular expression, allowing for longer loops (53, 54) beyond the usual 7 (15, 16), and returning longer sites wherever nested PQSs exist. The resulting sites were then filtered to include only the 619,282 PQSs that have a mapped experimental G4-seq data available, with no other such PQS within the 50 nucleotide (nt) range from its borders. The latter consideration was to make sure that the G4-seq measured mm% values, due to their lower resolution, would not reflect the cumulative scores of more than one G4 structure. Each such extended PQS from that database was then assigned a single mm% value reflective of its G4 structure formation propensity and stability (52). The assignment of the mm% value was done by selecting the highest one of all the 15-nt binned mm% data measured in G4-seq experiment (4) that overlaps with the PQS and its 50-nt flanks. This scheme was adopted in order to capture the peaks of the relatively broader G4-seq signals stemming from the respective PQS site.

### Processing of the UV damage data

The genome-wide maps of ultraviolet (UV) radiation-induced pyrimidine-pyrimidone (6-4) photoproduct (PP) and cyclobutane pyrimidine dimers (CPDs) (**Figure 2A**) were taken from (55), as accessed through the GEO (51) repository under the accession identifier GSE98025. We investigated the dataset of UV-exposed naked DNA, with data on the four major UV-damage types: TT^PP^, TC^PP^, TT^CPD^ and CT^CPD^, two for each of the PP and CPD couplings. All the four types were analysed separately. The data from two available technical replicates were then combined, due to the low coverage and the overall low duplication (4-11 %) of the exact damage sites in the two experiments of exposing the DNA to 20 J/m^2^ UVC radiation. The fewer duplicated cases were counted only once. This resulted in strand-assigned damage datasets for the human nuclear genome for TT^PP^, TC^PP^, TT^CPD^ and CT^CPD^ UV-caused cross links respectively.

**Figure 2.**
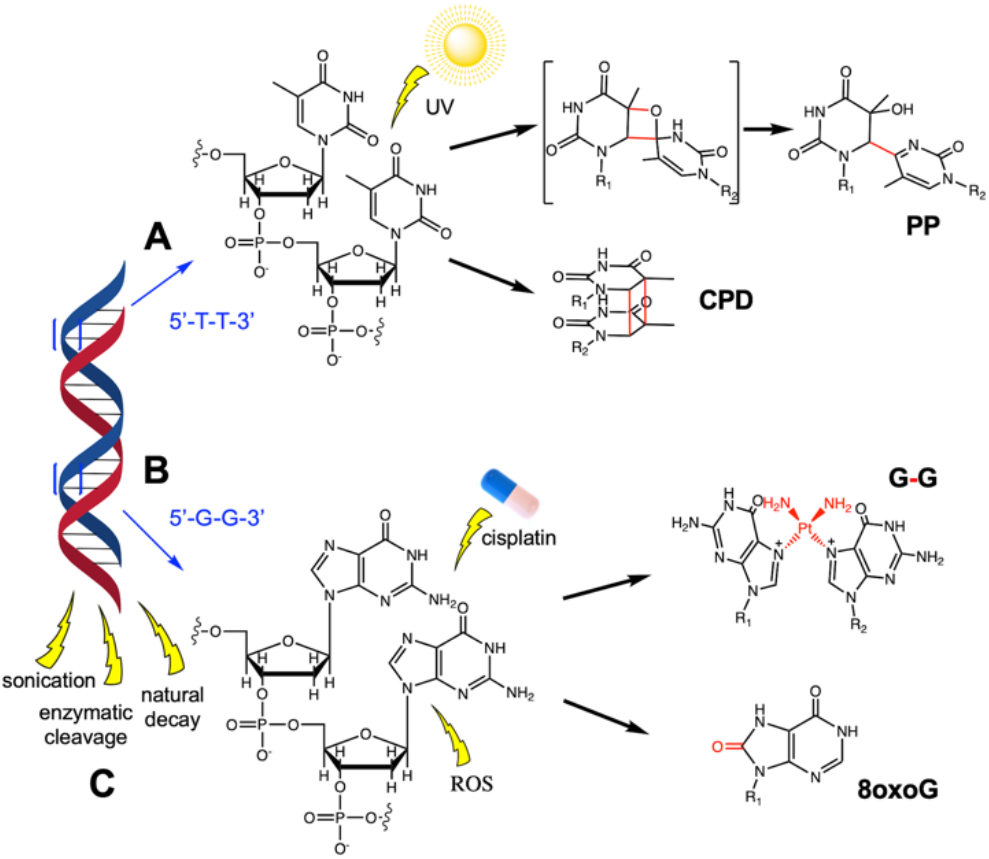
Chemical structures and schematic representation of DNA damage and fragmentation phenomena studied in this work. The example for ultraviolet (UV) radiation damage (**A**) is brought for TT pyrimidine dyad, forming two major types of cross-linking products, 6-4 photoproduct PP, shown along with the intermediate state, and cyclobutane pyrimidine dimer CPD. GG dyad site is shown in (**B**) to exemplify the formation of a cisplatin induced cross link, as well as the reactive oxygen species (ROS) caused oxidative product 8-oxoguanine (8oxoG). The deoxyribose phosphate residues are replaced with R_i_ substituents in each illustrated product, where the coupling and damage moieties are highlighted in red. Three other types of damage: natural decay-, artificial enzymatic- and sonication-induced DNA breakages are considered and highlighted in (**C**).

### Processing of the cisplatin damage data

The genome-wide maps of cisplatin cross link sites **(Figure 2B)** were taken from (56), as accessed through the GEO repository under the accession identifier GSE82213. We investigated the dataset of cisplatin-treated naked DNA leading to the prevalent G-G cross link product at GG dyad sites. The data from two available replicates were combined, resulting in a strand-assigned damage datasets for the human nuclear genome for cisplatin-mediated coupling between guanine residues.

### Processing of the 8-oxoguanine damage data

The genome-wide distribution of both 8-oxoguanine (8oxoG) oxidative DNA damage data **(Figure 2B)** were taken from (57), as accessed through the GEO repository under the accession identifier GSE181312. We investigated the dataset of 8oxoG-mapped naked genomic DNA, with the damages induced by KBrO_3_ treatment. The data from two available replicates were combined resulting in a strand-assigned 8oxoG damage dataset for the human nuclear genome.

### Processing of the DNA fragmentation data

The fragmentation sites were inferred by processing a number of primary sequencing data (**Figure 2C**). For the fragmentation *via* natural decay and fossilisation (58), the ∼80,000 years old Neanderthal genome BAM-format sequence alignment dataset from the Max Planck Institute for Evolutionary Anthropology was used, as accessed from the http://ftp.eva.mpg.de/neandertal/directory. For the enzymatically induced DNA fragmentation, the aligned fasta files were retrieved from NCBI Sequence Read Archive (SRA), deposited under the accession identifier SRX7808529 (59). For the ultrasound sonication-induced DNA fragmentation, we took the sequencing data available from The Simons Genome Diversity Project (60), randomly selecting the dataset with the NCBI SRA accession identifier ERR1019043. While the obtained datasets were already read-aligned, we ensured to take the end of the reads that mark the actual physical breakpoint, as opposed to the stop in sequencing pre-set by the sequencer as limited by the requested read size. We made sure that the identified sides of the aligned reads for the true breakpoints were indeed right and that the breakpoints we collected were of as high quality with as low of a background noise as possible, by cross mapping against both strands of the human hg19/GRCh37 reference genome and filtering out the ambiguous alignments.

### Overlapping damages with PQS sites of characterised G4 strength

To calculate the isolated effect of a G-quadruplex (G4) structure, separating it from the possible effects of underlying G-rich sequence, we used a strategy where the comparisons were only made between sequences that would all comply with the same overall PQS motif, whether making non-existent or weak to strong G4 structures. This way, while considering the damage *vs*. PQS crossover under the light of the gradual increase in the G4 strength, we would enrich for the effect of the structure from the possible sequence effects of the G-rich spans. PQSs were thus divided into eight categories based on their stability: PQSs with G4 stability score [0, 5], (5, 10], (10, 15], …, (35, 40] and >40 mm%. For each such stability category, we calculated the number of damage sites that had formed on the same strand as the PQS, and, separately, the number of damage sites formed on the strand opposite to the PQS (this was done separately for each damage type). The obtained counts were then converted into fractions from the number of available sites within the given (PQS or opposite) strand where the given damage type could in principle occur. The denominator in this fraction would thus depend on the exact damage type, i.e. number of all available TT sites for TT^PP^ or TT^CPD^ damage, number of available GG sites for cisplatin G-G cross links, number of available G sites for 8oxoG damage, length of the sequences for the NN breakpoints in DNA fragmentation. The latter fragmentation-caused breakpoints are not strand-specific, hence were analysed and plotted without strand differentiation. We next converted the described fractions (from all the damage types) to reflect their propensity change in % from the sequences with [0, 5] G4 stability score, which harbour PQSs with no G4 formation.

### DFT calculation of ionisation potentials of nucleobases and G-tetrad

Density functional theory (DFT) (61) calculations were done using KMLYP hybrid functional (62) with cc-pVTZ correlation consistent triple zeta basis set (63). The selected level of theory was shown to be the best out of a variety of DFT functionals for our purpose, with 0.73 eV average absolute error in predicting experimental ionisation potentials (IP) (64). The basis set employed for our systems is approximately equivalent to 6-31+G(d) in Pople’s definition, which is of appropriate complexity to obtain accurate results for IP and orbital energy calculations (65). All the calculations were done with prior geometry optimisation with (for a variant G and G-tetrad) and without symmetry constraints. The used nucleobases were all capped with hydrogen atoms, chosen as a replacement for the (deoxy)ribose phosphate moiety and to keep the molecular systems at a closed-shell configuration. The calculations were performed using Gaussian 03 suite of quantum chemistry programs (66). The KMLYP (62), inaccessible through a built in keyword, was achieved by requesting BLYP functional with iop(3/76=1000005570), iop(3/77=0000004430), and iop(3/78=0448010000) workflow modifiers suitable for only the Gaussian 03 input, as clarified on www.ccl.net exchange. The IP values were estimated from the highest occupied molecular orbital (HOMO) eigenvalues, in accordance with the Koopmans’ theorem IP = -E_HOMO_ (67), shown in multiple occasions to be of sufficient accuracy for most systems (64, 65, 68-70).

## RESULTS AND DISCUSSION

### Focus on structural effects at G4 sites on intrinsic damage

We considered the co-occurrence of various damage sites in our genomic DNA with sequence spans that have a prior potential to form G4 structures (**Figure 1**). Such putative quadruplex sequences (PQSs) were binned into categories according to their actual G4 structure formation mm% scores, as per the experimentally quantified (4, 52) values. This resulted in nine reporting categories for PQSs with G4 stability scores binned as [0, 5], (5, 10], (10, 15], …, (35, 40] and >40 mm%, where the G4 formation propensity rises from non-existent to extremely high, forming very strong G4 structures. This dimension (x-axes in **Figure 3**) thus enables us to have a unique differential view on the role of G4 structures, deconvoluting it from the mixed role of G-rich sequences that often fall under the PQS definition but may not form an actual G4. In other words, if we observe an effect that gradually changes with the gradual increase in G4 structure formation propensity, then the effect is likely to be structure driven. The used compendium of genomic damage types (**Figure 2**) with available datasets, were all analysed in this dimension, with the results normalised in accordance with the sites available to each type of damage. Therefore, the noted effects are normalised to a single damage site, hence the G4 effects on G damage or GG cross link will not be elevated by a mere accumulation of guanine bases in PQSs, and analogously, the effects of di-pyrimidine couplings caused by UV radiation will not be artificially depreciated because of the relatively low numbers of di-pyrimidine sites in PQSs (in the loops). Our revealed effects should thus be considered as of G4-caused structure (G4 and opposite single-stranded ssDNA, see **Figure 1A**-**C**) and of structure-derived electronic nature at G4 sites, acting on a single site available for a damage of a given type for each **A**-**F** plot in **Figure 3**. Additionally, since all our studied damage types, except DNA fragmentation breakpoints, are strand specific (**Figure 3A-D**), we can reflect on the structural effects of both G4s and the opposite strands that spend more time as ssDNA. The latter, exposed strand, effect is revealed when the damage is considered while happening in the strand opposite to the G4-forming one.

**Figure 3.**
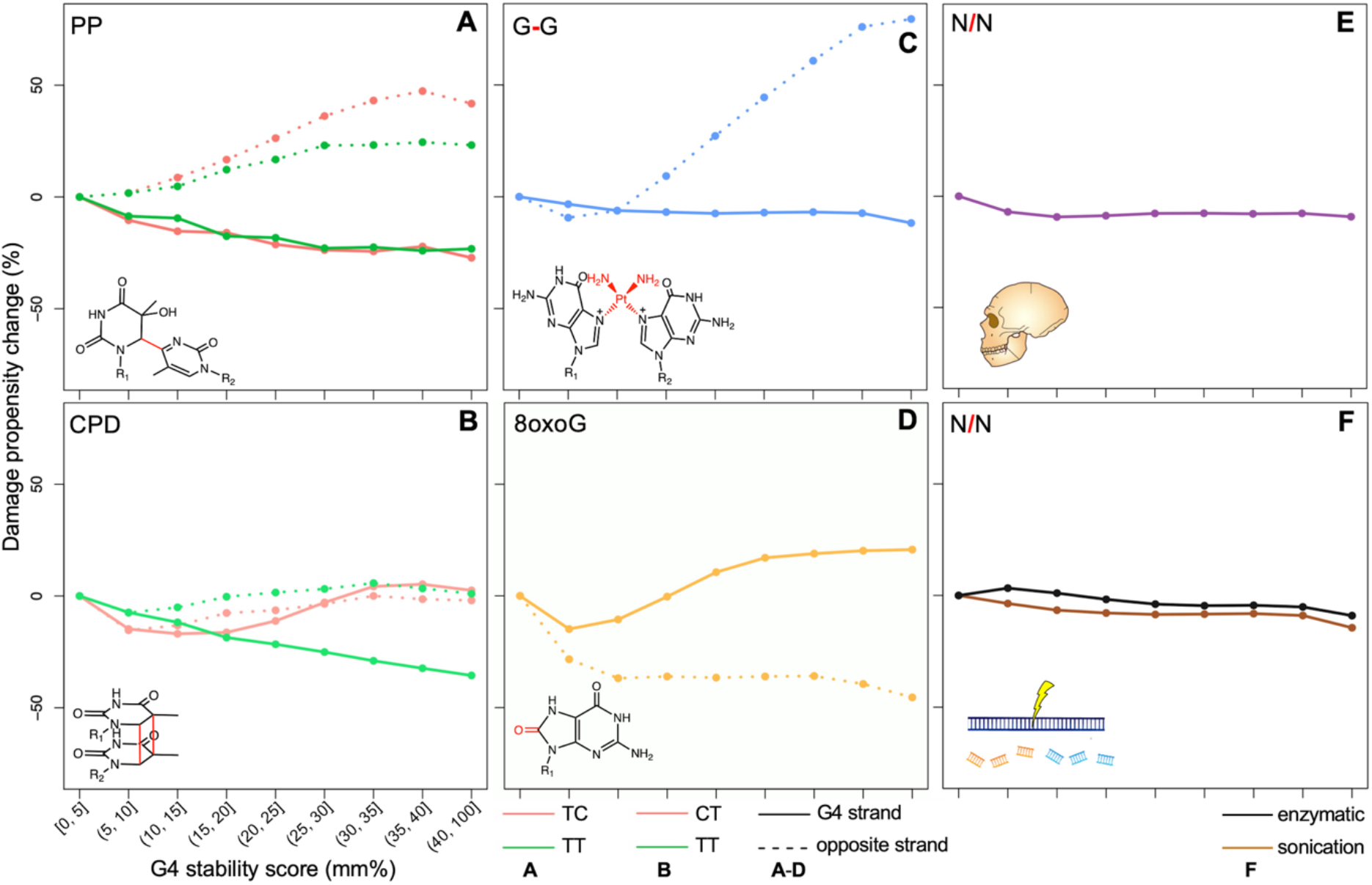
The differential damage propensity change in the sites of genomic putative quadruplex sequences (PQSs) with increasing G4 structure formation. The G-quadruplex stability (x-axes in **A**-**F**) was assessed through the G4-seq derived mismatch percentage (mm%) score, binned into ranges indicated at the bottom left plot. The damage propensity change (y-axes in **A**-**F**) is expressed as a percentage difference from the fraction of damage sites in PQSs with [0, 5] mm% negligible G4 formation, still conforming the used extended PQS motif [G_3+_N_1−12_]_3+_G_3+_. Therefore, all the plots start from 0 % for the damage propensity change at [0, 5] mm%. The prior fractions of damage sites were calculated as fractions of factually damaged sites from sites in principle available for a given damage type. Plots are shown for the ultraviolet radiation caused PP (**A**, for TC and TT dyads coloured in red and green respectively) and CPD (**B**, for CT and TT dyads coloured in red and green respectively) cross links, cisplatin-mediated G-G cross links (**C**), oxidative 8-oxoguanine (8oxoG) damage (**D**), breakage propensity change in between any two N bases from ancient DNA as inferred from fragmented Neanderthal genome (**E**), and breakage propensity change during artificial fragmentation *via* ultrasound sonication (brown) and enzymatic (black) treatment (**F**). Since the plots **A**-**D** represent strand-specific damage phenomena, the results are shown for both G4 containing strand (solid line) and the opposite strand (dashed line), which can both be “+” or “-”, depending on a specific PQS site. The dashed lines for the opposite strand in the plots **A**-**D** represent the changes in damage propensity while the strand spends increasingly more time in a single-stranded state, in front of increasingly more stable G4 structure.

### UV-caused damage at pyrimidine dyads

Exposed to UV radiation, DNA can become damaged where photochemical reactions allow for two adjacent pyrimidine base pairs, prevalently at TpT, TpC, and CpT dyads, to cross link together, forming a pyrimidine dimer (71). Such lesions, if not repaired, may then become the reason for mutations and UV-caused skin cancers (72). There are two major classes of UV-induced DNA lesions (**Figure 2A**): pyrimidine-pyrimidone (6-4) photoproduct (PP), and cyclobutane pyrimidine dimers (CPDs). PPs occur when a bond is formed between positions 6 and 4 of adjacent pyrimidines. PPs are mostly prevalent at TpT and TpC sites (55, 73, 74). CPDs are formed between the 5, 6 double bonds between two adjacent pyrimidines, usually more common between TpT and CpT sites (55, 74). In the context of G4 formation (**Figure 1**), even though the guanine-rich G4 strand contains less pyrimidine dyads prone to UV damage (**Figure 1B**), mostly located in loop regions, the opposite strand (**Figure 1C**) would be more enriched in such dyads involving cytosine. However, in any case, our results are all normalised by the counts of the damageable sites for each considered four (TT^PP^, TC^PP^, TT^CPD^ and CT^CPD^) damage types, and reflect on the G4 structure-driven effects in the G4 sites on a single damageable dyad moiety, not biased by the overall counts of such moieties in those loci. The results are summarised below, and show significant structure-driven effects on intrinsic UV-damage susceptibility.

#### UV-caused PP damage

For the UV-caused coupling into the PP damage product, we found that G4s tend to show a protective effect (a ∼25 % reduction of PP damage susceptibility at any single damageable site) on the strand in which they form (**Figure 3A**). This is reflected by the negative damage propensity change as the G4 structural stability increases in **Figure 3A**. Interestingly, the extent of the G4 protective effect is invariant to the specific pyrimidine dyad type, being very similar for TT^PP^ and TC^PP^ damages. On contrary, the damageable sites residing in the opposite strand become more vulnerable to UV damage, where we saw UV damage at the sites of more stable G4s increase by ∼50 % for TC^PP^, and ∼25 % for TT^PP^. It is worth noting that the PP damage susceptibility level is plateauing at the highest most extreme mm% ranges, likely due to the saturation in G4 stability. The overall trends observed for the PP damage are logical and typical to G4 sites: G4 structures form a tight knot on the G4 strand (**Figure 1A** and **B**), hence decreasing the expected level of UV exposure and post-exposure damage, but leave the opposite strand exposed to the UV radiation (**Figure 1C**) and highly dynamic to undergo the PP coupling. However, as can be seen from the below subsections, the most expected trends do not always stay true for the other damage types.

#### UV-caused CPD damage

The structural effects at G4 sites on CPD couplings are substantially different (**Figure 3B**) from that of PPs. In the CPD case, the increase in G4 stability, hence G4 formation propensity, seems to first protect both strands against CPD formation, then, for stronger G4s, damagability levels reach the values very close to the original ones at the negligible-G4 category ([0, 5] mm%). This trend is true for both G4 and opposite strands, and for both TT^CPD^ and CT^CPD^ damage products (**Figure 3B**). The only exception is the TT^CPD^ coupling at the G4 forming strand, where a stable G-quadruplex structure plays a protective role decreasing the damage propensity at a TT site in the loop regions by up to ∼35 % (green solid line in **Figure 3B**). The overall absence of substantial increase in CPD damage vulnerability, even while the pyrimidine dyads reside in the exposed opposite strand, could still be explained through a structural effect. Our observation is likely due to the key difference between the spatial arrangements of the two pyrimidines that facilitate the CPD coupling reaction, as contrasted to the PP coupling. There is a strict stacked arrangement of individual pyrimidines in the CPD product (**Figure 2A**). That spatial alignment between adjacent pyrimidines is already found in the normal Watson-Crick base-paired double helical DNA, even without the CPD cross links. To illustrate this, **Figure 4A** highlights the arrangement of two pyrimidines (T bases in blue), with the 5, 6 double bonds favourably positioned for the cycloaddition reaction towards CPD coupling while the DNA is in a double helical form. To this end, CPDs are less likely to form in the loosely structured and/or single-stranded DNA opposite to G4 structures, or within the loops of G4s. In weak G4s, the presence of misalignments in such transient folds likely prevents the formation of CPDs, hence explaining the decrease in CPD formation at (in both the same and opposite strand of) unstable G4s. The subsequent increase in UV-damage formation for more stable G4s can likely be attributed to the reduction in transience of G4 structures, resulting in more fixed loop structures where some stacking between two pyrimidines for CT, but not TT, sites may still be possible, thus cancelling out the overall protective effect.

**Figure 4.**
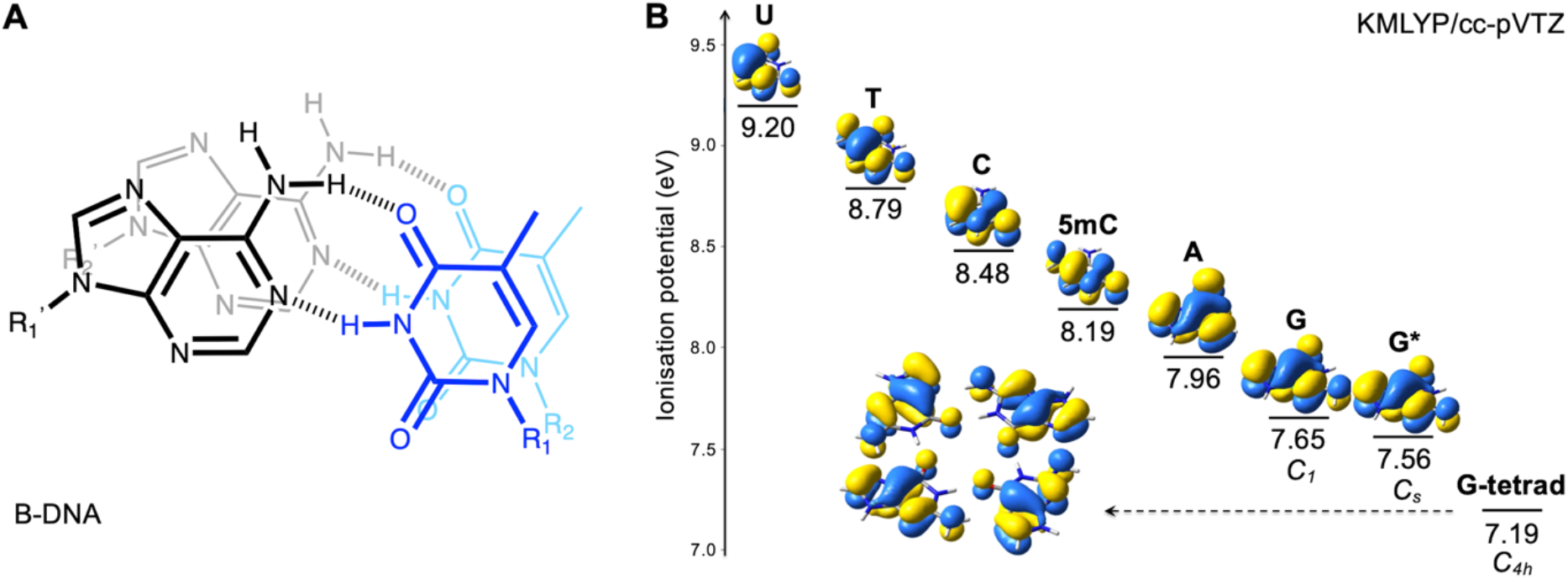
Structural and electronic considerations for the ultraviolet (UV) radiation-caused and oxidative damages. (**A**) shows the schematic view of the arrangement of neighbouring pyrimidines, relevant for UV damage, in double helical B-DNA. The example brings two Watson-Crick paired A:T base pairs adjacent to each other, hence formed by the moieties ApA and TpT in the respective two strands. The view is from top; the darker base pair is the one closer to the viewer; hydrogen bonds are shown *via* dashed lines; deoxyribose and phosphate residues are replaced with R_i_ substituents at each base. The comparative diagram (**B**) shows the calculated ionisation potentials for all DNA and RNA bases (A, T, G, C, U), the most common epigenetic modification 5-methyl cytosine (5mC), and the planar G-tetrad structure as a building block of G4 structures. The DFT calculations were done *via* KMLYP hybrid functional with cc-pVTZ basis set. The calculations for guanine were done both unconstrained (*C*_*1*_) and under *C*_*s*_ symmetry constraint, as indicated on the diagram. The calculation for the G-tetrad was performed under *C*_*4h*_ symmetry constraint. The molecular illustrations show the highest occupied molecular orbitals for each calculated system.

### Cisplatin cross links at GG sites

Cisplatin is the most widely used chemotherapy agent which acts by cross linking guanine bases in DNA and leading to cell death (75). The advent of recent sequencing technologies to map the cisplatin binding sites (56, 76), revealed a preferential coupling happening at GG dyads, with a non-random distribution leading to characteristic gene expression response (77, 78) and mutational signature (79) in cisplatin treated tissues. Here we analysed a single-nucleotide resolution cisplatin binding data from (56), which prevailed in cross-linked GG dyad sites.

The G4-site structural effects on the cisplatin-mediated coupling between guanine residues in GG dyads are summarised in **Figure 3C**. In this case, for the G4 strand, we have very little protection from the G4 structures on individual GG couplings within G-quadruplexes, with the damage propensity decreasing by ∼5-10 % (solid line in **Figure 3C**). Please note again, that the effect is normalised to a single damageable site, hence the overall summed propensity would be higher because of the presence of multiple GG sites on the G4 strand. On contrary, the effects on the damageability in the opposite strand is one of the most pronounced observed for the cisplatin-induced coupling (dashed line in **Figure 3C**), reaching up to ∼80 % increase in damage vulnerability at a single GG site residing on the opposite strand of a strong G4, as compared to opposite strands of PQSs with negligible G4 formation. Interestingly, it was noted before that mitochondrial genome is the major accumulation target for cisplatin cross links (76). Interestingly, eukaryotic mitochondrial genome is notorious for its overall GC-skew (80), featuring a G-, hence G4-, rich strand and C-rich opposite strand. Our observation can thus explain why we have so much cisplatin targeting of mitochondria, as the individual G susceptibilities in cisplatin mediated coupling will not change much in the G-rich strand, though with the elevated numbers of GGs multiplying their quantity, while the fewer GGs in the opposite strand will possess the noted substantially increased vulnerability to cisplatin cross links.

### 8-Oxoguanine damage of guanines

Reactive oxygen species, inevitably forming and circulating in a living matter, cause endogenous oxidative damage in our DNA (81). One of the most prevalent such damage is the oxidative modification of guanine nucleobase. Most commonly it forms the 8-oxoguanine (8oxoG) product, which has important implications for mutations, disease, and ageing (81-83). Despite the widespread presence of G nucleobases interspersed throughout the human genome, significantly deviating regional variations were noted for 8oxoG damage and repair (84), well beyond the expected G-rich telomers (82) and the noted spatial positioning in a nucleus (85). Prior low, megabase resolution mapping of 8oxoG in a genome revealed enrichment at around preferential recombination and single-nucleotide polymorphism sites (86), with a later work showing the role of 8oxoG in spontaneous mutations in mice (87).

Guanine is most vulnerable to oxidative damage due to its lowest ionisation potential (IP) of all nucleobases (69, 70, 88, 89). The low IP ensures easier loss of an electron, hence oxidation. Early works showed the increased propensity to ROS damage at the 5’ G within GG dyads (90, 91) due to further reduction of the IP in such systems, and the IP shown to be reduced even further when there are more consecutive adjacent Gs in a stacked formation in DNA (13). The IP reduction is the consequence of a strong delocalisation of highest occupied molecular orbital (HOMO) across the stacked G:C floors in DNA, also shown to contribute into coherent charge transport in GC-rich sequences (92). Our own density functional theory calculations, performed in this work, show that a well-formed G4-tetrad (**Figure 1A**) has even lower IP of 7.19 eV (see **Figure 4B**, brought in comparison to the other nucleobases), from 7.56 eV for an isolated G. The IP can potentially become even lower in a stacked G-tetrad arrangement within G4 structures (**Figure 1B**), thus potentially contributing to the increase in propensity to 8oxoG damage in G4s.

Recently, there has been a number of important contributions by Fleming and Burrows, demonstrating the possibility of fine interplay between the ROS damage at non-B DNA structures in modulating gene regulation, while investigating the consequence of 8oxoG formation at G4 and Z-DNA sites (13, 14). With the very recent publication of the seminal work by Hu and co-workers (57), first time depositing a nucleotide resolution genomic profile of 8oxoG damage for human, here we have a chance to investigate the purified and strength-differential structural effects at G4 sites on the predisposition to this type of damage, by operating on human genome G4 and 8oxoG datasets, all measured for naked genomic DNA.

8oxoG damage brings an example of a structure-driven electronic effect that deviates once more from the other observations from all our investigated damage types (**Figure 3D**). Unlike the UV-caused PP (**Figure 3A**) and cisplatin damage (**Figure 3C**), here we can see a reversal of the trends for the G4 and opposite strands. For 8oxoG damage, the opposite to G4 strand is always protective for the engulfed Gs (dashed line in **Figure 3D**), decreasing the propensity of a single G to be damaged by up to ∼50 % if facing the most stable G4s. This observation goes in hand with the electronic studies, as such strands will spend more time in a less structured and highly dynamic ssDNA state, hence minimising the possibility of HOMO orbital delocalisation and IP reduction present at well-formed standard B-DNA conformations. The G4 strand (initial section of the solid line in **Figure 3D**) behaves similarly protective to the engulfed Gs for the weak G4s, regardless of the presence of multiple Gs which were supposed to lower the IP hence the barrier to oxidation. Similarly, the explanation is likely due to the breakaway from standard B-DNA conformation, but still not forming strong and stable G4s. Overall, the above observations agree with the general mapping of 8oxoG sites at around PQSs, showing the surprising outcome of having lower propensity to 8oxoG damage in G4s (57). However, in our design, we could look at the differential effects of G4 structures at varying G4 stability levels. Owing to this, we can see that the Gs residing in increasingly more stable G4 structures with mm% above 20 (see the corresponding section of the solid line in **Figure 3D**), show a reversal of the noted trend and become moderately more vulnerable to 8oxoG damage, with the damageability at a single G site increasing by up to ∼22 % while residing in the most stable G4s. This reversal thus captures the structure-driven electronic effect (IP reduction) noted above for the tracks of Gs and G-tetrads (**Figure 4B**). However, the reflection is very marginal, which can mean that such electronic effects depend a lot on the stability of the structure and the absence of too much dynamicism around the stacked and co-planar molecular arrangements necessary for the delocalisation effects to stay intact. In fact, we even noted an increase of IP while allowing a negligible out-of-plane distortions in G, by going from C_s_ to lower C_1_ symmetry (**Figure 4B**). The essential rigidity necessary for the manifestation of the decreased IP and increased 8oxoG damage, most probably is possible in well-formed B-DNAs, and seems is overall hard to achieve in G4 structures.

### Natural DNA fragmentation upon decay and fossilisation

One of the challenges of studying ancient DNA from fossilised remnants is that such DNA is highly fragmented (93, 94). With the environmental factors playing important roles in the preservation of DNA in ancient species, a study found that the fragmentation is more correlated with thermal fluctuations rather than the sample age (95). Here, we analysed the fragmentation pattern from the Neanderthal genome (58) from the ∼80,000 years old remnant, as mapped on human genome considering their extremely high similarity (96, 97) and the expected lower number of recombination sites across the two species relative to the vast amount of fragmentation sites upon decay and fossilisation. Since the breakpoints are not strand specific, the analysis was done with no strand consideration, mapping the breakpoints that are for both “+” and “-” strands, on any G4 site on either “+” or “-” strand. The results show no significant differential effect from G4 structures (**Figure 3E**), with only very small, ∼5 %, protection at any single NN site residing in the highest G4 stability categories. This observation can either be the reflection of the exposed opposite strand and well packed G4 strand effects cancelling each other in terms of the breakage predisposition, effectively resulting in little effects from G4 sites of varying strength, or can simply be due to the over-exposure to fragmentation in such datasets, with too many short reads covering most of the sites with little bias.

### Artificial DNA fragmentation

#### Ultrasound sonication

DNA library preparation is required before the DNA can be sequenced on next-generation sequencing (NGS) platforms. This is accomplished by DNA fragmentation into short reads in high quantities. A commonly used method to fragment the DNA is *via* ultrasound sonication, in which the input DNA sample is subjected to high-energy ultrasonic waves that mechanically shear the DNA into fragments of varying size. Works emerged comparing and linking the sonication-induced sequence-dependent cleavage in DNA with other commonly used fragmentation methods (98), local flexibility of B-DNA (99), or chromatin accessibility in cells (100). The result of our G4 analysis for strand-invariant breakpoints induced by ultrasound sonication is shown as a brown solid line in **Figure 3F**. There is only a negligible protective effect (up to ∼13 %) from the G4 strength on any NN site with the breakpoint happening in between.

#### Enzymatic fragmentation

Restriction enzymes are used as an alternative cost-effective fragmentation method, though producing a larger number of sequencing artefacts and biases (101-103). The differential G4 analysis, however, resulted into a profile (see the black solid line in **Figure 3F**) very similar to ultrasound sonication result but with less protective effect (up to ∼7 %). Overall, for artificial fragmentation procedures, whether through sonication or enzymatically, the two possible explanation brought for the natural fragmentation may still hold true.

### General notes

In summary, here we demonstrated an approach that enables us to enrich for and isolate the structural effects from intra-strand G-quadruplexes and opposing ssDNA on various DNA damage types in genomic G-rich regions. The outcomes demonstrate that, in such regions, the structural effect does not consist of a simple “shielded G4” and “exposed-opposite-strand” duality, but rather can have different significant manifestations for various damage types, and can even vary depending on the G4 strength. We also exemplified a damage type, 8oxoG, where the structural changes influence damageability through the alterations and coherence in electronic delocalisation effects, shedding light on and unifying the seemingly contradictory results in past considerations (13, 57). Overall, our results outline the multifaceted nature of genomic G4 sites, and enable the demonstration of the linked structure-driven effects on 9 types (including 4 subtypes of UV damage cross links and 3 types of strand-invariant DNA fragmentation) of DNA damage phenomena. Considering the immense biological relevance of the G4 sites (2, 3, 19) and their specific distribution in our genome (4, 21, 25, 28, 33, 40), the differential influence of these loci on DNA damage can thus be an important non-specific driver of an emergent complexity in genome organisation and gene regulation. With intrinsic sequence effects on DNA damage and mutation acting at below 10 bp range (104, 105), here we also establish a structure-mediated meso-scale (∼30 bp range on average) sequence effects on DNA damage susceptibilities, showcasing the different modes and magnitude of the effects depending on the damage type, relative strand localisation and G4 strength.

## DATA AVAILABILITY

The computer code, necessary to reveal the effect of G-quadruplex structures on any supplied type of genomic DNA damage, can be accessed through the following GitHub repository: http://github.com/SahakyanLab/G4Damage. All the used public datasets are accessible from the established genomic data repositories as detailed in **Materials and Methods**.

## ACKNOWLEDGEMENTS

The Sahakyan Laboratory has been supported by the UK Medical Research Council (MRC), MRC Strategic Alliance Funding (MC_UU_12025). AA is grateful to the Kazanah Foundation for the studentship supporting his DPhil studies, CF is grateful to the University of Cambridge Trinity College Rouse Ball Research Fund for supporting her MPhil internship in Oxford. PP is grateful to MRC, Hertford College, Clarendon Fund and Radcliffe Department of Medicine for supporting his DPhil studies.

